# Improved Analysis of Phage ImmunoPrecipitation Sequencing (PhIP-Seq) Data Using a Z-score Algorithm

**DOI:** 10.1101/285916

**Authors:** Tiezheng Yuan, Divya Mohan, Uri Laserson, Ingo Ruczinski, Alan N. Baer, H. Benjamin Larman

**Affiliations:** Department of Pathology, Division of Immunology, Johns Hopkins University School of Medicine, Baltimore, MD 21205, USA.; Department of Genetics and Genomic Sciences, Icahn School of Medicine at Mount Sinai, New York, NY 10029, USA.; Department of Biostatistics, Bloomberg School of Public Health, Johns Hopkins University, Baltimore, MD 21205, USA.; Division of Rheumatology, Department of Medicine, Johns Hopkins University School of Medicine, Baltimore, MD 21224, USA.

## Abstract

Phage ImmunoPrecipitation Sequencing (PhIP-Seq) is a massively multiplexed, phage-display based methodology for analyzing antibody binding specificities, with several advantages over existing techniques, including the uniformity and completeness of proteomic libraries, as well as high sample throughput and low cost. Data generated by the PhIP-Seq assay are unique in many ways. The only published analytical approach for these data suffers from important limitations. Here, we propose a new statistical framework with several improvements. Using a set of replicate mock immunoprecipitations (negative controls lacking antibody input) to generate background binding distributions, we establish a statistical model to quantify antibody-dependent changes in phage clone abundance. Our approach incorporates robust regression of experimental samples against the mock IPs as a means to calculate the expected phage clone abundance, and provides a generalized model for calculating each clone’s expected abundance-associated standard deviation. In terms of bias removal and detection sensitivity, we demonstrate that this z-score algorithm outperforms the previous approach. Further, in a large cohort of autoantibody-defined Sjögren’s Syndrome (SS) patient sera, PhIP-Seq robustly identified Ro52, Ro60, and SSB/La as known autoantigens associated with SS. In an effort to identify novel SS-specific binding specificities, SS z-scores were compared with z-scores obtained by screening Ro-positive sera from patients with systemic lupus erythematosus (SLE). This analysis did not yield any commonly targeted SS-specific autoantigens, suggesting that if they exist at all, their epitopes are likely to be discontinuous or post-translationally modified. In summary, we have developed an improved algorithm for PhIP-Seq data analysis, which was validated using a large set of sera with clinically characterized autoantibodies. This z-score approach will substantially improve the ability of PhIP-Seq to detect and interpret antibody binding specificities. The associated Python code is freely available for download here: https://github.com/LarmanLab/PhIP-Seq-Analyzer.

## Introduction

The binding specificities of circulating immunoglobulins harbor a wealth of information related to the exposure history and health status of vertebrate organisms. Unbiased techniques to extract this information provide new opportunities for linking environmental exposures to disease, and discovering antigenic determinants of complex immune-related pathologies, including autoimmunity, allergic response, and malignancy. Several complementary high-throughput platforms have been developed to address the challenges of antibody profiling.(1–4) We have previously described the development of Phage ImmunoPrecipitation Sequencing (PhIP-Seq), which is based on T7 bacteriophage display of synthetic oligonucleotide encoded peptidome libraries. Antibody-peptide binding is quantified via high-throughput DNA sequencing of the immunoprecipitated phage libraries (5–9).

The PhIP-Seq method provides several key advantages over alternative approaches. First, synthetic DNA libraries can be made to cover complete or partial proteomes, including meta-genomically defined proteins and protein variants that do not exist in nature. This database-driven design feature was utilized for the construction and screening of the phage displayed human virome (‘VirScan’), which was accompanied by simultaneous high resolution alanine scanning of several dominant epitopes.(9) Further, the streamlined assay protocol takes place in solution phase (versus array-based solid phase techniques), requires no specialized equipment or instrumentation, and can be easily automated on standard liquid handling robots.(6) Perhaps the most important advantages of PhIP-Seq, however, derive from the use of high-throughput DNA sequencing technology as the assay readout. Sequencing of phage libraries is used to quantify clonal abundance with linear behavior over a large dynamic range. Critical to scalability, sample multiplexing via DNA barcoding enables screening at a per sample cost that is roughly one hundred fold less expensive than comparable array-based techniques.

Sequence reads, generated by deep sequencing of phage displayed peptidome libraries, are first demultiplexed and then aligned to the reference library sequences. For simplicity, we use exact sequence matching, which is computationally efficient and captures the vast majority of reads that arise from unmutated library members. Post-alignment read count data is then subjected to enrichment analysis, in order to detect peptide-antibody binding interactions (or peptide-bait interactions more generally). An ideal enrichment analysis approach should (i) account for inherent bias in the starting phage library as well as any bias introduced throughout the assay, (ii) appropriately incorporate the technical variation inherent to sequencing-based measurements, and (iii) consider characteristics of the data which are specific to the PhIP-Seq assay.

Our previous PhIP-Seq analytical strategy implemented a Generalized Poisson (GP) model to estimate the *p*-values associated with peptide enrichments. (5,6,9,10) Briefly, the clonal abundance distribution of the starting library was determined via sequencing. For intervals of abundance in the starting library, the observed abundance distribution of the corresponding peptides in an enriched library was fit to a GP distribution. Model parameters were estimated for each interval of the starting library. These parameters were regressed against the intervals of abundance, so that regressed model parameters could be used to formulate an expected GP distribution for all peptides, which varied smoothly over the entire range of starting abundance. This approach suffers from two main limitations. First, it does not consider the bias in the enriched library due to antibody-independent binding to the capture matrix. In the present work we demonstrate that this bias is significant. The GP distributions used to fit these biased populations are therefore overdispersed, compared to a model that uses matrix-alone enriched libraries to define the ‘expected’ distribution. The GP approach therefore produces both false positives (peptides that bind the capture matrix independent of antibody) and false negatives (peptides that are truly enriched but do not meet significance due to overdispersion of the model). The second limitation relates to quantification of measurement uncertainty. In the GP model, a given peptide’s enrichment significance is a function of the variance observed among all peptides of similar starting abundance. Preferably, however, enrichment significance should instead be a function of each peptide’s measurement variability, as determined by repeated measurement. Here we propose to address these limitations by developing a z-score enrichment algorithm that is based upon analysis of replicate negative controls (mock IPs).

In a typical PhIP-Seq experiment, we perform immunoprecipitations containing patient antibodies, as well as multiple negative control immunoprecipitations that lack antibody (i.e. mock IPs). Any capture matrix, including protein A/G coated magnetic beads, will exhibit some degree of background binding to the phage library, while the binding of specific peptide-displaying clones may be further enhanced (resulting in false positives). Ideally then, antibody-bound libraries captured on protein A/G beads should therefore be compared to mock IPs (i.e. libraries captured on protein A/G beads alone). In addition, measurement uncertainty can theoretically be quantified by analysis of these same negative controls if they are performed in replicate. By comparing antibody-enriched libraries to replicate negative controls, one can thus quantify increased abundance versus expected abundance, as well as the expected uncertainty in the measurement of each clone. With these parameters, enrichments can thus be reported as z-scores, a common and intuitive measure familiar to biomedical researchers (‘how many standard deviations away from the background value’). Rigorously, z-scores only apply to normally distributed continuous variables (rather than sequencing-generated discrete count data). In our view, however, the facile interpretation of z-scores balances this caveat. Compared to the GP-based method, the z-score approach should therefore increase the sensitivity and specificity of enrichment detection, as well as the interpretability of PhIP-Seq data.

In order to assess the performance of the z-score algorithm, we screened a collection of Sjögren’s Syndrome (SS) patients’ sera against a 90 amino acid (90-aa) peptide library spanning the human proteome.(10) SS is a systemic autoimmune disease, characterized by inflammation of the lacrimal and salivary glands with resultant impairment of tear and saliva production, and dryness of ocular and oral mucosal membranes. It may occur alone or in association with a second systemic rheumatic disease, such as systemic lupus erythematosus (SLE), rheumatoid arthritis, or systemic sclerosis. Antibodies to the Ro/SSA ribonucleoprotein complex (with specificity for the Ro52/TRIM21 and/or Ro60/TROVE2 antigens) are present in 60 - 80 % of SS patients, but are not specific for the disease, being also found in SLE and other rheumatic diseases, although at lower frequency. Antibodies to SSB/La (hereafter referred to as SSB) are also characteristic of the disease, but are present at lower prevalence and are almost always present in association with anti-Ro/SSA antibodies.(11) The presence of anti-Ro/SSA antibodies is a major criterion in current classification schema for SS, which are designed to ensure uniformity of disease definition for clinical trials or other research studies.(12) In the absence of anti-Ro/SSA and anti-SSB antibodies, the diagnosis of SS can only be established with a labial gland biopsy showing a characteristic salivary gland histopathology (i.e. focal lymphocytic sialadenitis). We therefore sought to detect autoantibodies more specific to SS, particularly for individuals lacking Ro/SSB autoantibodies.

## Methods

### Patient cohorts and autoantibody testing

The Johns Hopkins University Institutional Review Boards approved the collection of clinical data, serum and other biospecimens from patients for these studies. All patients were >18 years old and gave informed consent.

#### Sjögren’s syndrome

Sera were collected from 193 patients with Sjögren’s syndrome seen in the Johns Hopkins Jerome L. Greene Sjögren’s Syndrome Center between October 2008 and September 2015. All subjects met classification criteria for SS, 190 by the 2002 American-European consensus group criteria and 3 by the 2012 American College of Rheumatology criteria.(13,14) The predominant clinical phenotype in all patients was that of SS; however, some had overlap features with other systemic rheumatic diseases, including 10 with a non-erosive inflammatory arthritis and anti-CCP antibodies and 4 with limited systemic sclerosis and anti-centromere antibodies.

#### Systemic lupus erythematosus

The Hopkins Lupus Cohort is a prospective cohort in which patients with SLE are followed at least quarterly. Patient inclusion in the cohort is based on the clinical diagnosis of SLE by a member of the Rheumatology Division; 94% of the patients satisfied at least four of the 1982 American College of Rheumatology revised criteria for the classification of SLE.(15,16) For each patient, basic demographic characteristics, presenting and cumulative clinical manifestations, and immunologic markers have been recorded since cohort entry.

#### Autoantibody testing

Serum from each of the 193 patients was tested in the Johns Hopkins Rheumatic Disease Research Core Center laboratory for the presence of anti-Ro52, anti-Ro60, and anti-La antibodies. Antibodies against Ro52 and SSB/La were assayed using commercially available ELISA kits, per the manufacturer’s protocol (QUANTA Lite, Inova Diagnostics). Values below 20U were called “negative”, values between 20U and 79U were called “positive”, and values greater than 80U were called “positive(+)”. Ro60 antibodies were determined by immunoprecipitation of ^35^S-methionine-labeled Ro60 generated by *in vitro* transcription and translation, as previously described.(17) Each of the antibody assays was performed on the same patient serum sample as used for the PhIP-seq assay. Two out of the 193 patients were missing anti-Ro52, anti-Ro60 or SSB antibody clinical data.

### PhIP-Seq screening

PhIP-Seq screening was performed as described previously.(5,6,18) Briefly, we employed a mid-copy T7 bacteriophage display library spanning the human proteome, which consists of 259,345 90-aa peptide tiles that overlap adjacent tiles by 45-aa. After expanding the library to high titer using BLT5403 *E. coli* (Novagen), it was aliquoted and stored at −80 °C in 10% DMSO. Prior to immunoprecipitation, the IgG concentration of each serum sample was determined using an in house ELISA assay (capture and detection antibodies from Southern Biotech, catalog numbers 2040-01 and 2042-05, respectively). Antibody binding reactions occurred in 1 ml mixture containing 2.5 × 10^10^ particle forming unit of the human peptidome library (diluted in PBS) and 2 μg of IgG (typically ~0.2 μl of serum). After rotating the antibody binding reaction overnight at 4 °C, 20 μ! of protein A coated magnetic beads and 20 μl of protein G coated beads (Invitrogen catalog numbers 10002D and 10004D) were added to each reaction and rotated for another 4 hours at 4 °C. A Bravo (Agilent) liquid handling robot performed three bead washes, and then resuspended the beads in 20 μl of a PCR master mix containing Herculase II Polymerase (Agilent catalog number 600679). After 20 cycles of PCR, 2 μl of this reaction was added to a second 20 cycle PCR reaction, for the addition sample specific barcodes and the P5/P7 Illumina sequencing adapters. Sequencing was performed on an Illumina HiSeq 2500 in rapid mode (50 cycles, single end reads).

### Z-score algorithm

After sample demultiplexing, we obtained about 0.7 - 2 million reads per sample, which were then aligned to the reference library sequences using the bowtie aligner.(19) For each sample *j*, the resulting read count of each peptide *i* (RC_*ij*_) was normalized by total read counts for the sample (T_j_), then multiplied by one million. Read counts per million (RPM_*j*_) given a peptide at a sample is thus defined

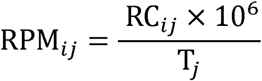

Calculation of pep-z scores of enrichment in an antibody-containing (‘experimental’) IP was performed as follows. 261 mock IPs across multiple PhIP-Seq experimental batches served as our negative control set. A cubic polynomial regression of the logarithm (base 10) of the standard deviations (*D_0_*) versus the logarithm (base 10) of the median RPM_*ij*_’s (*X_0_*) was fitted (Figure 2):

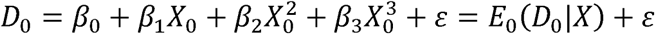

The ordinary least square (OLS) coefficients *β_0_, β_1_, β_2_*, and *β_3_* were estimated using the Python module *statsmodels*. This model was henceforth used to calculate the expected standard deviation for each peptide, given its RPM_*j*_.

For a single PhIP-Seq experiment, a set of (typically eight) mock IPs served as batch-specific negative controls. For each sample *j*, we implemented a robust linear regression of the RPM_*j*_’s (*Y_i_*) versus the mean of the mock IPs’ RPM_*j*_’s (*X_i_*) (using the RLM function of Python model *statsmodels*, where the Huber loss function was used for M estimation).

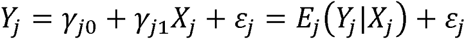

The expected values of *Y_j_*, determined by *E_j_*(*Y_J_|X_j_, γ_j0_, γ_j1_*), were used as sample-specific background. The corresponding deviations vector (*D_j_*) given *E_j_* was determined by the function *E_0_*(D_0_|*E_j_, β_0_, β_1_, β_2_, β_3_*). Therefore, the peptides’ z-scores vector for sample *j* is:

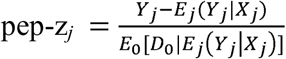

Under the null hypothesis of a two-sided normal distribution, all positive pep-z scores could be converted into *p*-values where appropriate. At the protein level, for *k* peptides associated with a given protein, promax-z scores are calculated from the set of pep-z scores as follows:

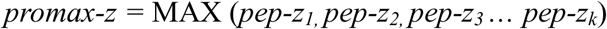

### Bioinformatics Workflow

PhIP-Seq data analysis by the PhIP-Seq Analyzer takes place in two stages. First, the *bioTreatEASTQ.py* script is run. This script demultiplexes fastq sequencing files, and build a sample information file, and sets parameters for running the pipeline. Second, the main script *bioPHIPseq.py* is run. This pipeline will launch the sequencing alignments, reads counting, data normalization and z-score statistics, sequencing quality control, polynomial regression, enrichment analysis, *etc*. Sequence alignment utilizes the Bowtie aligner (version1) with default parameters *-a –best –strata -l 40 -v O – norc –nomaqround –sam-nohead*. Only unique aligned reads were counted per peptide per sample, and a counting table with peptides in rows and samples in columns is constructed. Afterwards, tables are constructed, including RPMs, pep-z and promax-z scores tables with proteins defining the rows instead of peptides.

### Availability and requirements of the pipeline

The associated Python code is freely available for download at https://github.com/LarmanLab/PhIP- Seq-Analyzer. The software user manual including its installation and the required running environments is also incorporated in the package. We developed PhIP-Seq Analyzer using Python (version 3.4.0) programming language on a Linux operating system. The development and testing environment was LinuxMint 17.2 (X_86 64 bits) on a computer equipped with one Intel Xeon E5-1603 2.80GHz and 32 GB memory. The pipeline was also tested on the Maryland Advanced Research Computing Center (MARCC) computing cluster, running CentOS release 6.7.

## Results

### Accounting for library bias due to capture matrix binding

In previous reports, we assumed a uniform background binding of the phage library to the capture matrix (i.e. perfect correlation with the input library).(5,9) However, capture matrices may introduce substantial library bias, which we quantified with a series of matrix only (no antibody input) PhIP-Seq mock immunoprecipitations (“mock IPs”) using the human peptidome (10) and a mix of protein A/G coated magnetic beads (Figure 1A). At a Benjamini-Hochberg (BH) adjusted *p*-value ≤ 0.05, 313 peptides were differentially abundant after immunoprecipitation, compared with the unenriched input library (n = 10; Mann-Whitney-Wilcoxon test). Furthermore, the background binding of the library to the capture matrix was highly reproducible, as shown in Figure 1B. These data suggest that sample IPs are more appropriately compared to mock IPs, versus comparison to the unenriched starting library.

**Figure 1.**
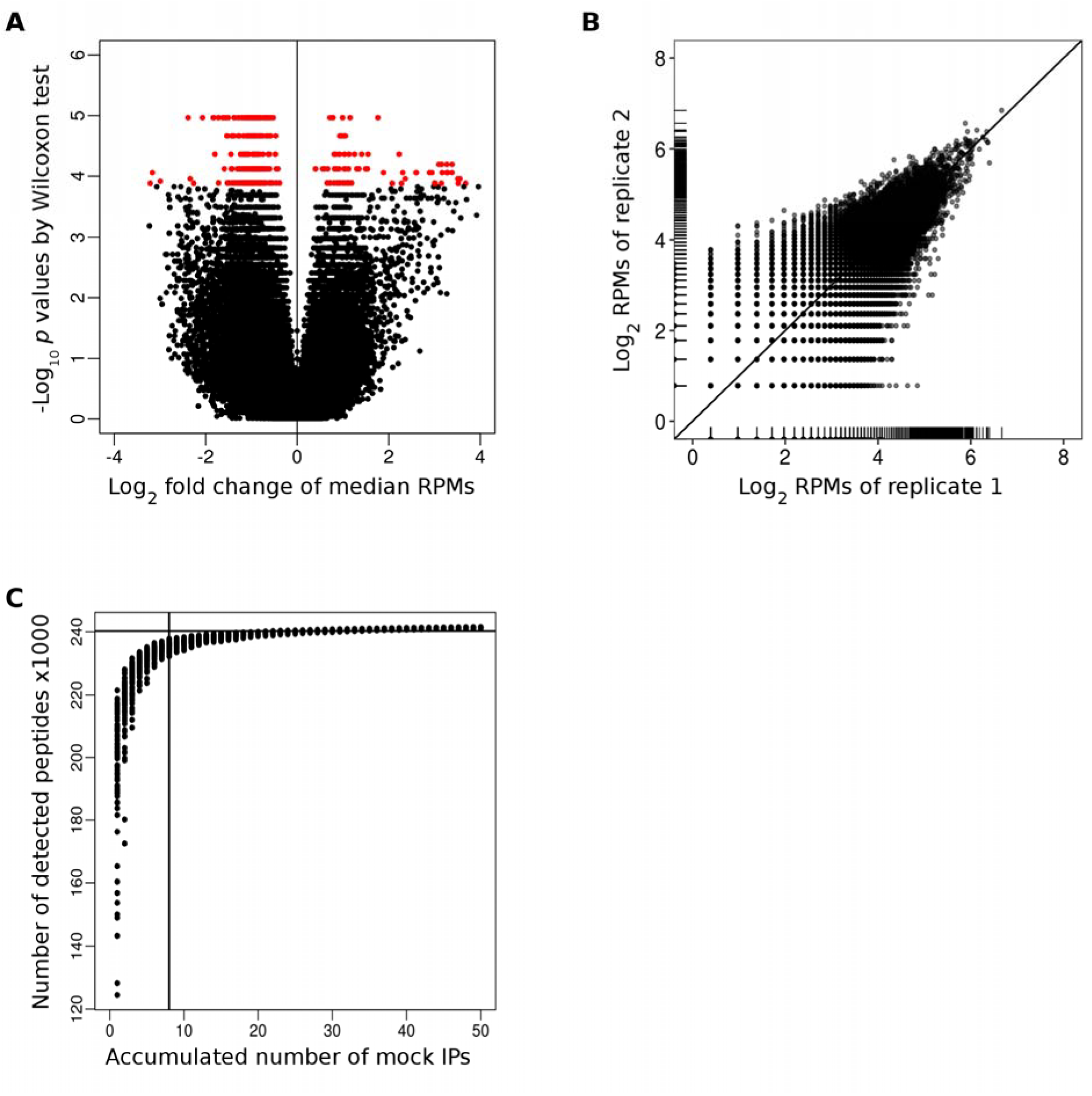
Accounting for matrix binding bias. **(A)** Volcano plot showing fold change of the median peptide RPMs of the mock IP versus those of the unenriched input library. Differentially enriched peptides identified by Wilcoxon test (BH-adjusted *p*-value ≤ 0.05) are shown as red dots. **(B)** Comparison of each peptide’s RPM from duplicate mock IPs. **(C)** Saturation analysis of mock IPs. A range of 1-50 mock IPs were randomly selected from 261 mock IPs, and the number of peptides detected was counted. For each number of mock IPs, the process was repeated and plotted 100 times. The horizontal line marks the number of peptides detected by sequencing the unenriched input library. The recommended minimum number of mock IPs is ~8 (vertical line) when using the 90-aa human library and protein A/G beads as capture matrix.

Though mock IPs better accounted for binding artifacts, they may fail to adequately sample the library due to bottlenecking of phage particles’ binding to the capture the matrix. In a mock IP, each library member’s representation will depend on the total complexity of the library, population skewness, and the strength of the binding to the capture matrix. To address loss of representation, the data from multiple replicate mock IPs can be aggregated. In this study, protein A/G coated magnetic beads were used for serum IgG capture experiments; we observed sampling saturation on this matrix with roughly eight mock IPs (Figure 1C). For 96 well plates of protein A/G IPs, we therefore included eight mock IPs per plate. We suggest performing this type of saturation analysis for each new library and capture matrix employed. An important benefit of performing replicate negative control IPs is that technical variation can also be measured and incorporated into peptide enrichment analyses, as discussed below.

### Development of z-score enrichment metrics for PhIP-Seq data

For biological measurements in which an expected ‘background’ value and standard deviation of measurement can be determined, the dimensionless ‘z-score’ (also referred to as ‘z-value’, ‘standard score’, or ‘normal score’) is commonly used to indicate the magnitude of a relative difference. Z-scores traditionally assume an underlying Gaussian distribution, which does not strictly apply to discrete PhIP-Seq read count data. We have nonetheless found this metric to be a useful and familiar concept for biomedical researchers wishing to interpret PhIP-Seq enrichment data. Here, we show how z-scores can be readily calculated by analysis of mock IPs.

For both mock IPs and experimental IPs, each peptide’s raw read count is normalized by the total read counts of the sample, and then multiplied by one million, to get RPM values. Expected sample-specific experimental RPMs in an antibody-containing PhIP-Seq immunoprecipitation are determined by linear regression of experimental RPMs versus the average RPMs observed on the mock IP. This approach is important because strong peptide enrichments ‘consume’ reads that would otherwise be available to the remaining, unenriched peptides. Without correcting for this ‘consumption bias’, the unenriched peptides would artifactually appear to be diminished compared to the mock IPs, and the magnitude of true enrichments would similarly be underestimated.(20,21)

We next sought to calculate the expected standard deviation of each peptide, given its expected RPM. To this end, we fitted a cubic polynomial to the logarithm of the standard deviations of the mock IPs’ RPMs plotted against the mock IPs’ median RPMs (Figure 2A). Use of median RPMs instead of mean RPMs for calculation of each peptide’s expected abundance in the mock IPs provides protection against accidental contamination of mock IPs’ with experimental IPs, spurious clone enrichments due to poor library preparations, and overly bottlenecked or undersequenced libraries.

**Figure 2.**
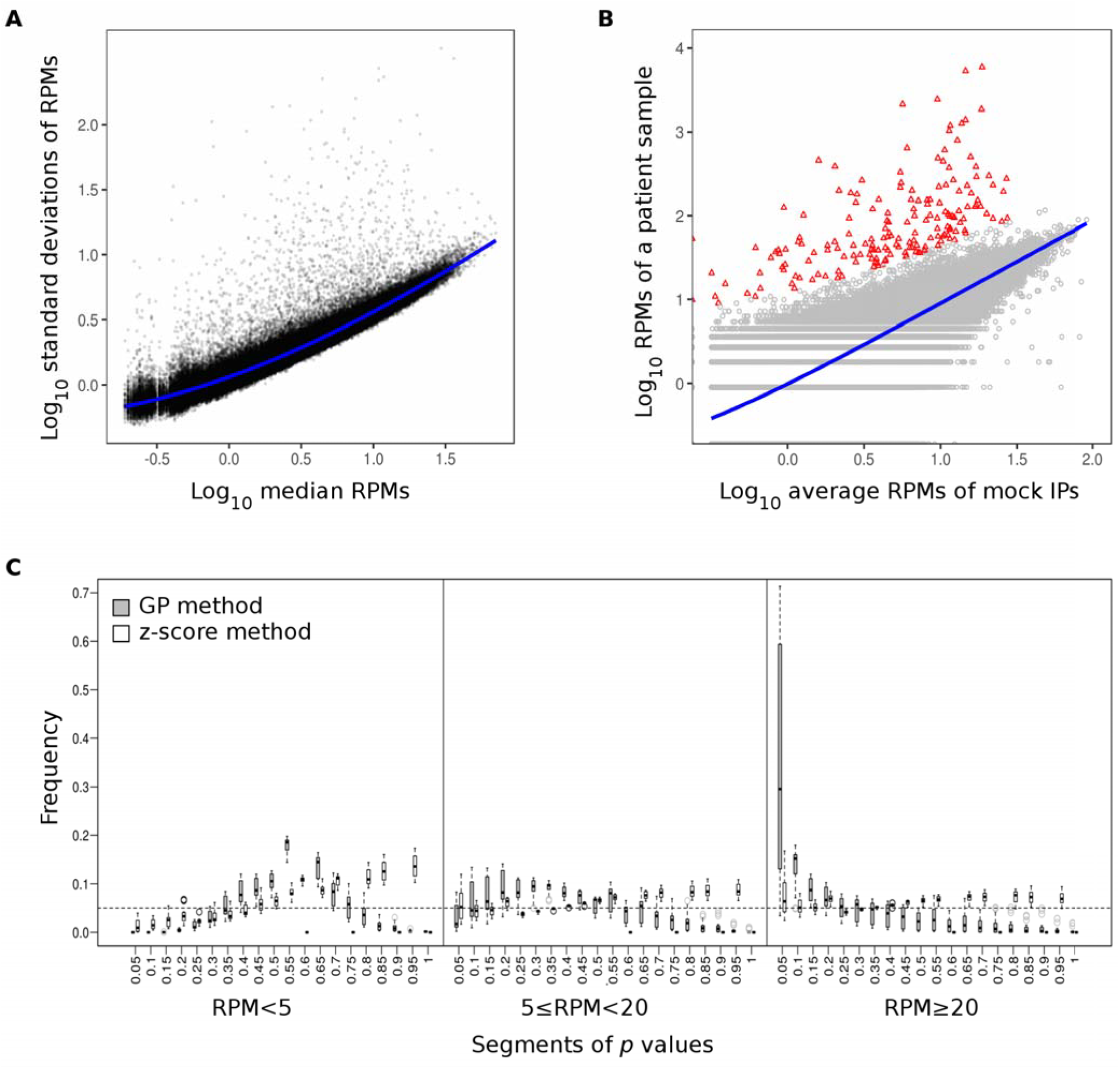
The z-score peptide enrichment metric. **(A)** Cubic polynomial regression (blue line) of standard deviations of peptides’ logged RPMs versus their logged median values among 261 mock IPs. **(B)** Robust linear regression (blue line) of RPMs from an experimental sample versus the mean RPMs of eight mock IPs from one 96 well PhIP-Seq experiment. Red triangles indicate peptides with pep-z scores above 10. **(C)** Histogram of *p*-values over the 0-1 interval, assessed at different sampling depths (n=15 mock IPs); comparing uniformity of the z-score method (open boxes) versus the GP method (gray boxes). The horizontal dotted line at 0.05 is shown to represent ideal fitting to the null model.

Each peptide’s experimental ‘pep-z score’ is then equal to the difference between the experimental RPM and the expected RPM, divided by the expected standard deviation, as showed in Figure 2B. Experimental samples are regressed against the mean or median of the within-batch mock IPs (e.g. from same 96-well plate). For peptides with zero expected read counts, their expected standard deviations are set equal to the minimum value of the regression model. The resulting pep-z scores can then be used to rank peptides, or to identify which peptides are enriched above a threshold (as in Figure 2B at a threshold of 10, colored red).

One important feature of any statistical model is the uniformity of the *p*-value distribution over the zero-to-one interval. Figure 2C shows the uniformity of *p*-value transformed z-scores for a set of 15 negative controls, and compares this to the *p*-value distribution obtained from the same data but analyzed instead using our previously published GP-based *p*-value estimator.(5,6) This analysis revealed that a more uniform distribution of *p*-values, especially for RPMs greater than 20, is achieved using the z-score based analysis, consistent with a more accurate null model of peptide binding. Importantly, the z-score approach tended to provide better protection against type I error, compared to the GP approach.

Next, we determined how false discovery rate (FDR) varied as a function of pep-z score threshold. For this analysis, we plotted the FDR versus z-score thresholds for 177 experimental IPs, assessed by comparison with the mock IPs from the same plate (Figure 3A). At an FDR of 0.05, the pep-z score threshold varied between 5 and 15, with a median value of ~10. In the current study, we therefore considered a pep-z score of 10 to be reliable. If enrichment scores are to be considered quantitative, the larger pep-z scores should be systematically associated with increased reproducibility. We assessed this property of the pep-z scores by determining the correlations of three sets of technical duplicate experimental IPs. Pep-z scores of peptides enriched in either duplicate exhibited predominantly tight correlations over at least two logs (Figure 3B). Importantly, the number of significant hits above the pep-z score threshold was also relatively consistent between duplicates (Figure 3C).

**Figure 3.**
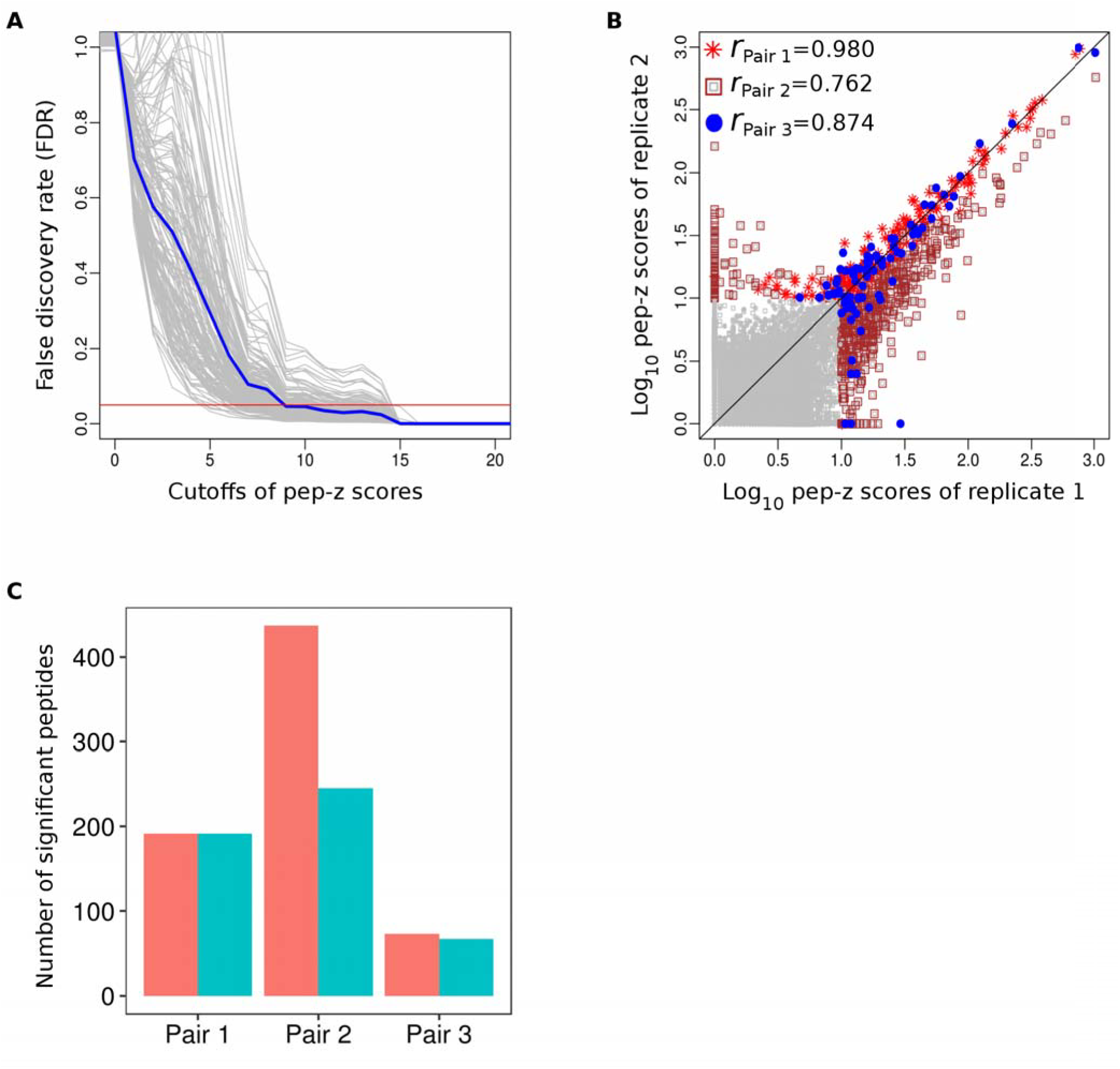
Detecting significantly enriched peptides. **(A)** False discovery rates (FDR) of significantly enriched peptides. FDR is the ratio of the numbers of significant peptide enrichments for each of 176 experimental IPs to the average number of peptide enrichments from 15 mock IPs, over the range of z-score thresholds shown. The blue line is the median FDRs at each pep-z score threshold. The intersection of the 10% FDR line (red horizontal line) with median FDR curve (blue line) occurs at a pep-z score threshold of ~10. **(B)** Reproducibility of pep-z scores across three independent sets of technical replicates. Correlation factors are provided at the top of the plot. Peptides above threshold (>10) in either duplicate are colored. **(C)** The number of peptides enriched above threshold are shown for each set of technical duplicates.

### Use of z-scores to characterize autoantibodies in SS patients

We next assessed the performance of PhIP-Seq in a comprehensive analysis of 193 SS patients’ autoantibodies using the 90-aa human peptidome library.(10) Each serum sample was clinically tested for the presence of the three most widely measured autoantibodies: anti-Ro52, anti-Ro60 and anti-SSB. The results of these routine clinical assays were considered to be ground truth in this study. In order to directly compare peptide z-scores with the corresponding clinical full-length protein based assays, we extracted the maximum pep-z from the set of peptides designed to represent each of these proteins (‘promax-z score’). Figure 4 compares the results of the clinical assays with the corresponding promax-z scores. At a promax-z score threshold of 10, PhIP-Seq status was roughly consistent with their clinical serostatus; PhIP-Seq sensitivity was the highest for sera strongly anti-Ro52 positive.

**Figure 4.**
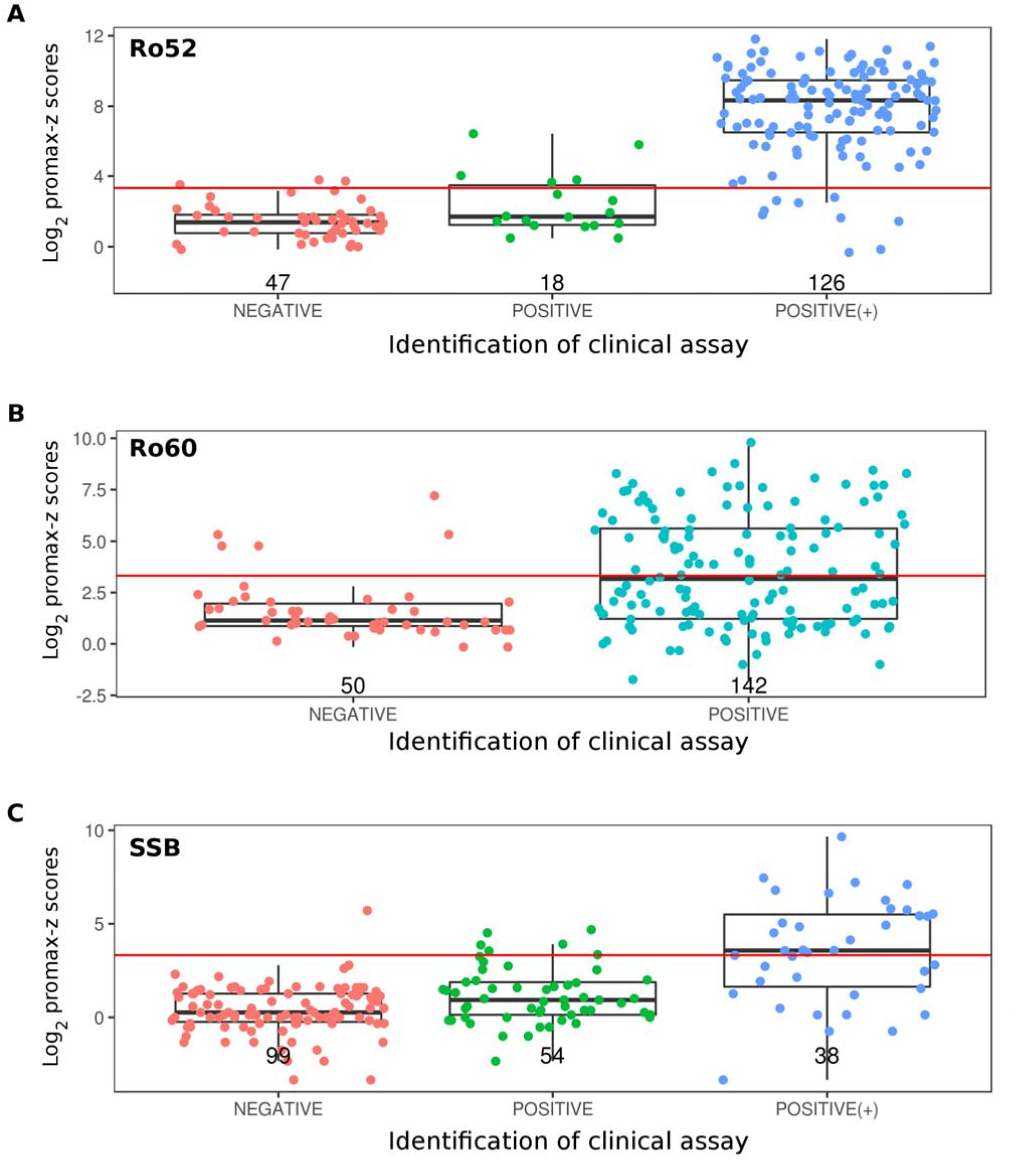
Comparison of PhIP-Seq z-scores with standard clinical assays. PhIP-Seq promax-z scores for Ro52 **(A)**, Ro60 **(B)**, and SSB **(C)** were compared against the results of the corresponding clinical diagnostic tests with the number of patients shown below. The promax-z score threshold (>10) is shown by horizontal lines.

Comparing Ro52 versus Ro60 promax-z scores revealed a pattern of exclusivity. Whereas anti-Ro52 antibodies were frequently detected in the absence of Ro60 antibodies (~27%, upper left quadrant of Figure 5), the converse was almost never the case (~1%, lower right quadrant of Figure 5). This pattern of fine specificities has been previously noted with other assay systems.(22–26) In contrast, however, anti-Ro60 antibodies were present in 12% of Ro52 negative patients by clinical assay.

**Figure 5.**
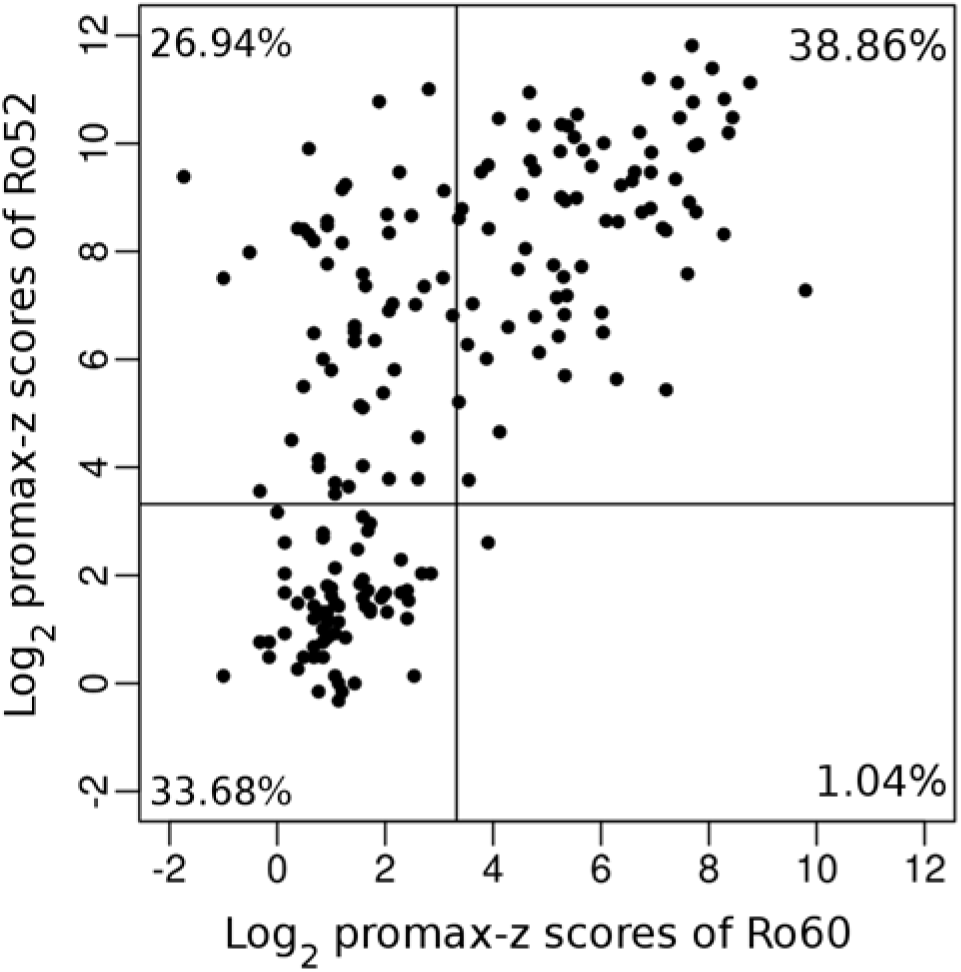
Ro52 versus Ro60 PhIP-Seq single positives. Whereas PhIP-Seq Ro52^+^/Ro60^−^ patients were frequent (27%), Ro52^−^/Ro60^+^ patients were rare (1%). Percent positives are labeled in each quadrant.

At the epitope level, PhIP-Seq revealed the most frequently targeted peptides from these dominant SS antigens; we identified at least three non-overlapping common epitopes within Ro52, at least one in Ro60, and at least two in SSB. To visualize these data, we clustered the patients based on their patterns of peptide enrichments for each of the three autoantigens. Interestingly, several samples classified as Ro60 negative by the clinical assay, clustered tightly with the true positives based on their PhIP-Seq peptide enrichment pattern (Figure 6). This result raises the possibility that the peptidome library presents cryptic epitopes not accessible to patient antibodies tested against the native protein. Comparison of protein-level and peptide-level analyses may therefore provide complementary disease-relevant insight, such as patterns of epitope spreading or antigen hierarchy, for example.

**Figure 6.**
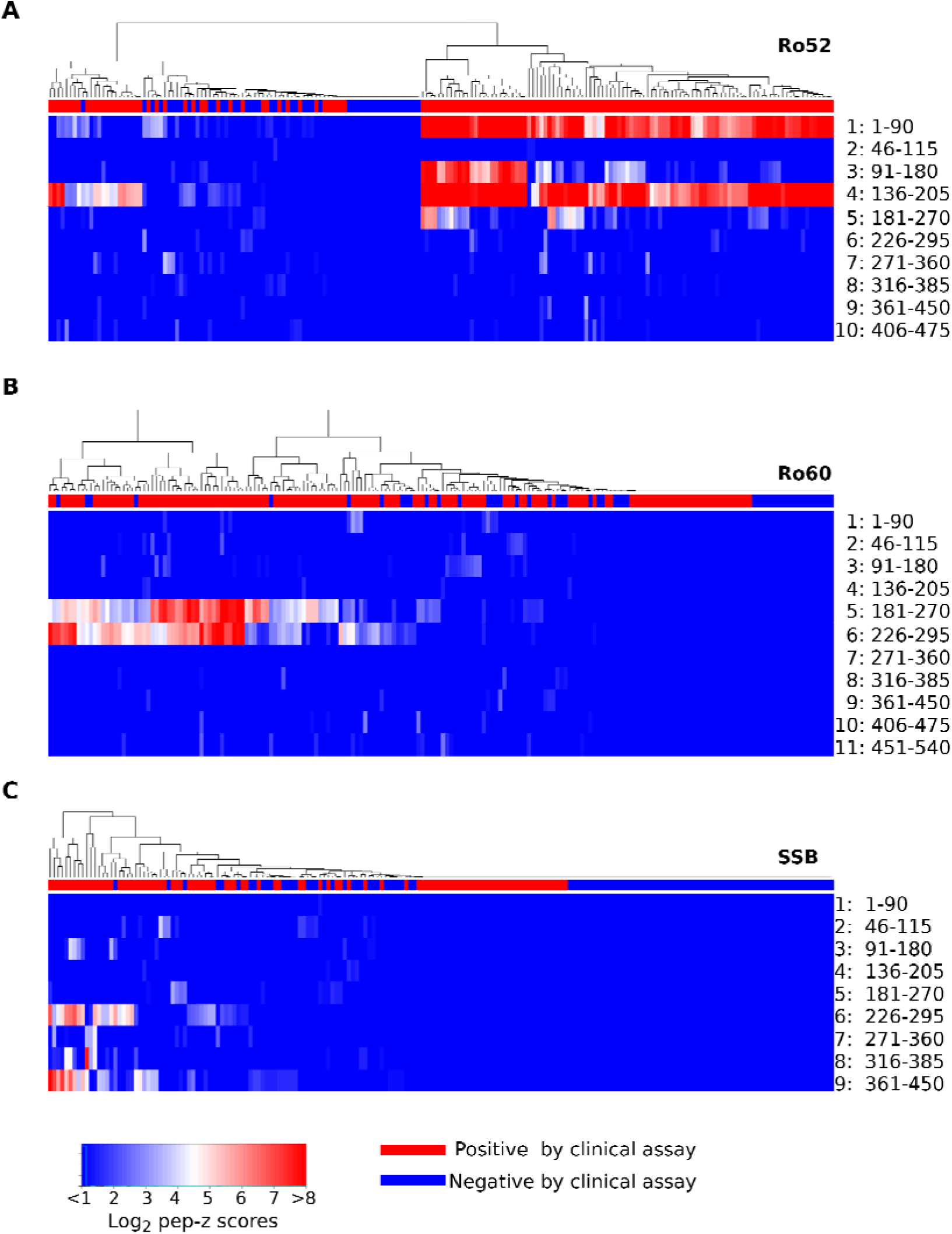
Epitope level analysis of SS antigens. Enrichment of each overlapping peptide are shown for each protein: Ro52 **(A)**, Ro60 **(B)**, and SSB **(C)**. The amino acid positions from N- to C-terminus (top to bottom) are labeled on the right. The density of gradient color bar shows log2 pep-z scores. Patients in columns identified as positive or negative by the corresponding clinical test (red or blue along the top bar) were clustered by pattern of peptide enrichment.

We next performed receiver operator characteristic (ROC) analyses on the SS data set, again assuming the clinical assays to be ground truth. PhIP-Seq achieved an AUC of 93.2%, 71.5% and 73.4% for Ro52, Ro60, and SSB using the z-score method, compared with an AUC 92.1%, 71.7%, and 69.0%, respectively, using the GP method (Figure 7). The z-score method therefore slightly improves discrimination between cases and controls using these three autoantibody biomarkers.

**Figure 7.**
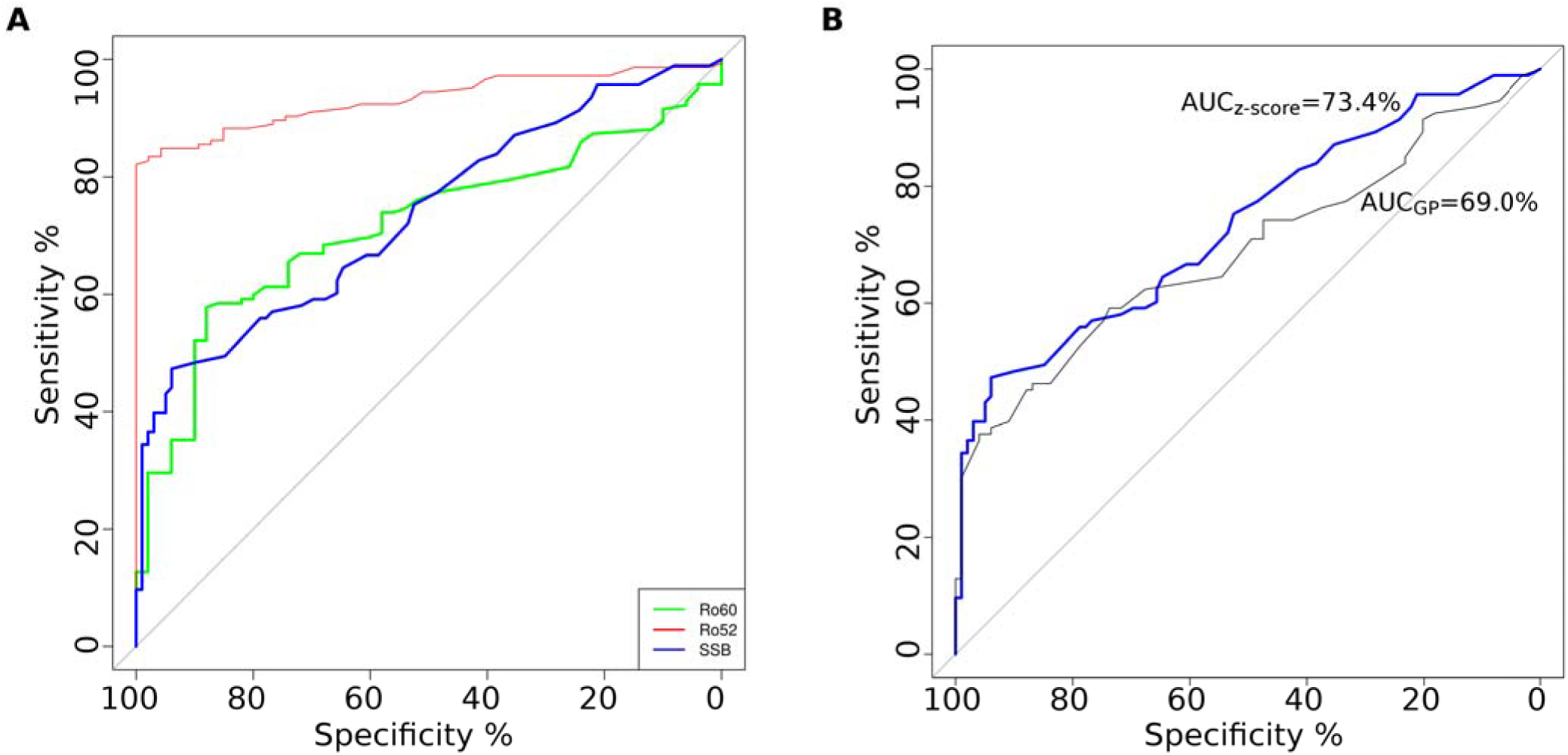
ROC comparison of z-score versus GP analysis. **(A)** ROC analysis using promax-z scores for Ro52, Ro60, and SSB considering the clinical test results to be ground truth. **(B)** ROC curves for SSB determined using z-scores (blue) or the GP statistic (gray).

Finally, we employed the z-score method to search for novel SS-specific autoantigens. We used Fisher’s exact test to calculate the significance of differential autoantibody frequency for each human protein among SS patients and a set of 301 non-SS controls. A volcano plot of this result is shown in Figure 8A. SSB, Enrichments of Ro52, Ro60, and SSB in SS patients were statistically significant (*p*-values < 10^−12^) by this analysis. SS is considered a systemic rheumatic disease, and as such shares Ro52, Ro60, and SSB autoantibody prevalence with other systemic rheumatic diseases, such as lupus. We therefore sought to identify SS-specific autoantibodies by comparison with autoantibodies in lupus patients who did not have SS, but who harbored Ro52 antibodies. In this analysis, neither SSB nor Ro60 were significant using Fisher’s exact test (Figure 8B). Several previously reported lupus-specific autoantibodies were identified in this comparison (not shown), but no unreported SS-specific autoantigens were identified. These results are in agreement with other antibody assay systems, including gel diffusion with extracts of human B-lymphocytes, (27,28) homogenates of calf thymus and human spleen, (29) as well as solid phase assays using purified or recombinant antigens, (30–33) suggesting that common SS-specific autoantibodies, if they exist, are most likely directed against discontinuous or post-translationally modified epitopes not present in our human peptidome library. Our results also confirm findings of Burbelo *et. al*. regarding the extraordinary antigenicity of the Ro52 antigen in SS, when studied with a luciferase immunoprecipitation system.(34)

**Figure 8.**
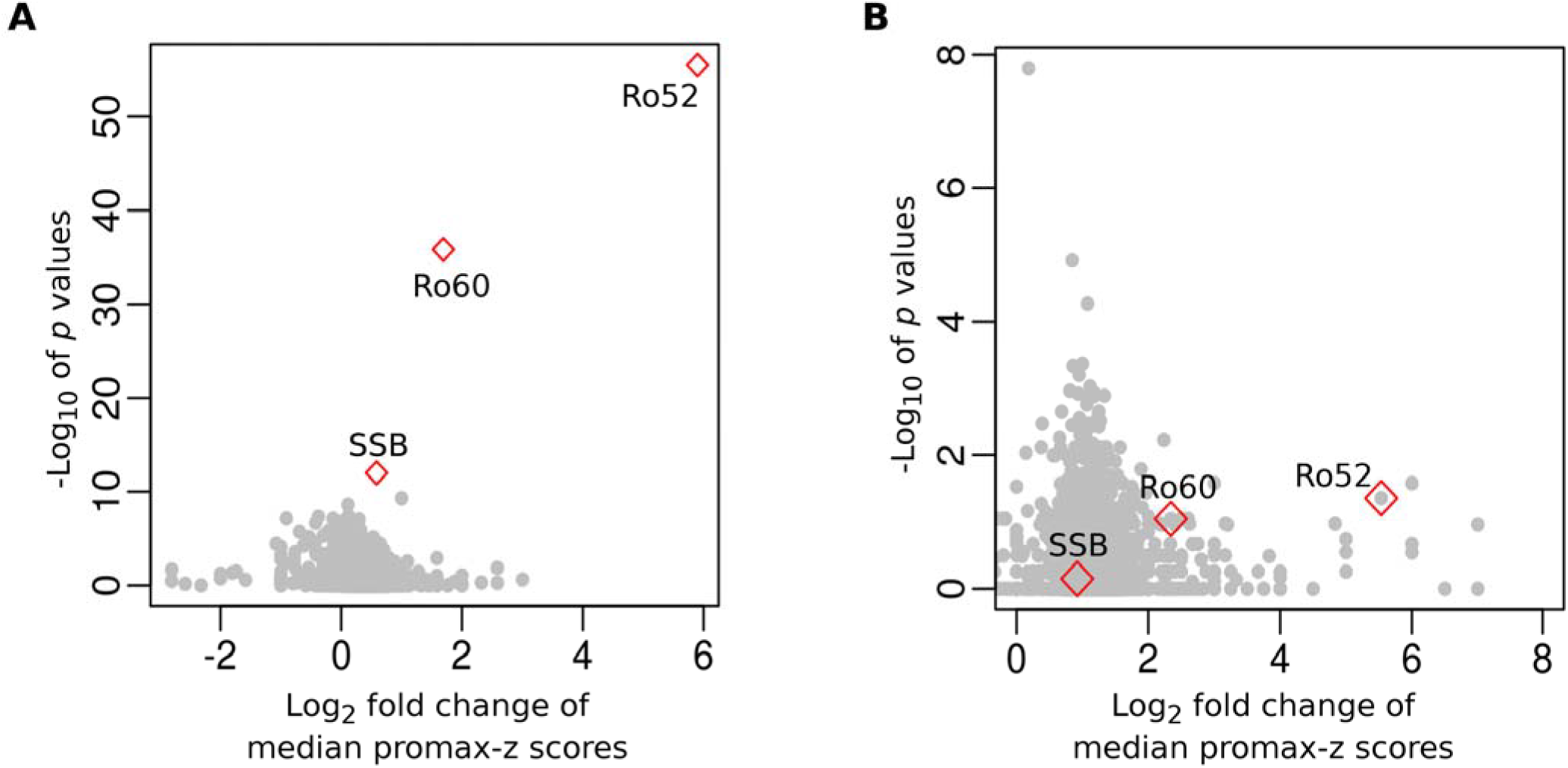
Proteome-wide analysis of SS autoantibodies. **(A)** All promax-z scores were assessed for disease association in an analysis of SS autoantibodies versus non-SS control autoantibodies. **(B)** Promax-z scores were then used to identify SS-specific autoantibodies by comparison with Ro52 positive lupus patient autoantibodies.

## Discussion

The PhIP-Seq z-score algorithm presented here provides several key advantages over our previously published generalized Poisson (GP) approach. It removes bias due to nonspecific binding of the capture matrix, exhibits increased sensitivity and specificity, and provides a measure of antibody-dependent peptide enrichment that is more intuitive to biomedical researchers. The z-score method also provides additional flexibility, in that sample-to-sample comparisons can similarly be made (instead of the sample-to-mock IP comparisons made here). Disadvantages associated with the z-score approach include the requirements for additional negative controls (mock IPs) and the construction of a model for the expected standard deviations specific to the capture matrix. Given the findings presented here, however, we conclude that the superior performance of the z-score algorithm outweighs these additional requirements.

Compared with more traditional antibody binding assays, such as ELISA or western blotting for instance, PhIP-Seq is an indirect measurement, and involves additional sources of potential artifacts. For example, analysis of the phage DNA requires PCR amplification, so that even small differences in per-cycle amplification efficiency may introduce nonlinear differences in the resulting normalized read counts for the same clone in two different samples. At the Illumina sequencing stage, differential clustering efficiency may also skew the final read counts. During sequence demultiplexing and alignment, stringency-defining parameters may impact the final data set in subtle and unpredictable ways. Perhaps most importantly, however, are the effects of clonal dropout, which can be due to either population bottlenecking or insufficient sequencing depth.

The inability to accurately estimate the abundance of undersampled clones makes quantifying their enrichment particularly challenging. Relatively undersampled clones will exhibit a high level of variability, and thus uncertainty; importantly, this uncertainty is naturally incorporated into the z-score metric. An uncertainty-incorporating metric is essential for comparing enrichments to one another, and thus ranking them in order of their likelihood for validation using a secondary assay. If however, certain peptides are *a priori* known to be enriched, the magnitude of their relative changes may more accurately be quantified using the fold change of their normalized read counts, rather than using their z-scores.

Discrepancies between PhIP-Seq and clinical assay results were most prominent for Ro60. The clinical assay, using immunoprecipitation of Ro60 antigen generated by in vitro translation and transcription (IP/IVTT), detected anti-Ro60 antibodies in a significantly higher number of sera than PhIP-Seq. This suggests the importance of conformational epitopes in this reactivity. With PhIP-Seq, anti-Ro60 antibodies were almost exclusively found with anti-Ro52 antibodies. They were present alone in ~1% of SS subjects when assayed by PhIP-Seq and in ~12% of SS subjects, when assayed by IP/IVTT. Ro52 and Ro60 are encoded by different genes, localize to different cell compartments and are not part of the same stable macromolecular complex.(35–38) Reactivity to the Ro60 has been previously shown to depend on conformational epitopes; reactivity is largely lost with denaturation of the protein.(39) In contrast, most sera recognize linear epitopes of the denatured Ro52 antigen, in agreement with the high rate of detection via PhIP-Seq.(38)

One important advantage of PhIP-Seq over full-length protein based assays is the determination of peptide level binding specificities. These data may provide disease relevant information, such as epitope-disease phenotype relationships,(40) or guide the development of improved diagnostic assays.(34) Screening SS sera, we determined the presence of multiple non-overlapping commonly targeted epitopes within Ro52, Ro60, and SSB. Furthermore, several patients who tested negative for Ro60 using a clinical assay were found to be convincingly positive by looking at the peptide-level PhIP-Seq z-scores, suggesting the presence of a cryptic epitope not available to certain patients’ antibodies in the form of a native protein. The most striking discordance between PhIP-Seq data and the clinical tests, however, is its high rate of false negativity, particularly for SSB (A total of 40 and 91 out of 193 SS patients were detected by PhIP-seq and ELISA, respectively.). This poor sensitivity may be the result of several limitations associated with PhIP-Seq. The dominant epitope(s) may be absent from the phage library due to their discontinuous nature, requirement for disulfide bond formation or other post translational modification, or the epitope displaying phage may happen to be missing or under-represented in the library. Alternatively, the targeted protein’s individual peptide enrichments may be weak (resulting in a low promax-z), whereas a more aggregated statistic may reveal a signal above background. Such metrics have been proposed for the analysis of RNA-Seq data but have not been explored here.

There are ways in which future analyses of PhIP-Seq data may be improved. For example, small differences among clones’ PCR amplification efficiency may be quantified using serial dilution of the input library. Computationally correcting for these differences may enable more accurate comparisons across samples and/or peptides. Experimental spike-ins may also prove useful for data calibration. For instance, addition of a fixed number of monoclonal phage particles to the first PCR reaction may facilitate estimation of particle numbers, particularly when PCR primers are depleted during amplification. One could also spike a monoclonal antibody into the immunoprecipitation reaction, which may improve calibration by accounting for differences in washing stringency, etc. These and other experimental modifications, designed to improve the robustness and/or sensitivity of the PhIP-Seq assay, can readily be incorporated into the z-score paradigm developed in this study. Ongoing development of the open source PhIP-Seq-Analyzer software package is therefore anticipated.

## Acknowledgements

We are very grateful to Michelle Petri and Wei Fu for providing serum samples from Ro-positive SLE patients without SS. Funding: This work was made possible by a Discovery Grant from the Jerome L. Greene Foundation, an NIH U24 award (AI118633), NIH R01 awards (CA194042, AI095068 and DE12354-15A1), a Prostate Cancer Foundation Young Investigator award, and a Catherine and Constantinos J. Limas Research Award.

## Author contributions

AB, TY and HBL conceived the project and wrote the manuscript. DM performed the PhIP-Seq assays. TY wrote the software and analyzed the PhIP-Seq data. UL and IR provided statistical support.

